# Assortative mating and within-spouse pair comparisons

**DOI:** 10.1101/2020.02.17.949347

**Authors:** Laurence J Howe, Thomas Battram, Tim T Morris, Fernando P Hartwig, Gibran Hemani, Neil M Davies, George Davey Smith

## Abstract

Comparisons between cohabitating spouses have been proposed as an aetiological design method to reduce confounding and evaluate effects of the shared adulthood environment. However, assortative mating, a phenomenon where individuals select phenotypically similar mates, could distort associations. We evaluated the use of spousal comparisons, as in the within-spouse pair (WSP) model, for aetiological epidemiological research.

Using directed acyclic graphs and simulations, we demonstrated that the WSP model can reduce confounding if spouses are correlated for an unmeasured confounder, but that WSP comparisons are susceptible to collider bias induced by assortative mating. Empirical analyses using spouse pairs in UK Biobank found evidence that genetic association estimates from the WSP model are attenuated compared to random pairs for single nucleotide polymorphisms (SNPs) associated with height (shrinkage: 23%; 95% CI 20%, 25%), educational attainment (74%; 95% CI 66%, 81%) and body mass index (23%; 95% CI 14%, 32%) as well as for an alcohol consumption SNP (29%, 95% CI 5%, 46%). Some of these attenuations are likely to reflect effects of assortative mating because height and educational attainment are unlikely to be strongly influenced by the adulthood environment. In contrast, effect estimates of increasing age on coronary artery disease and systolic blood pressure were found to be concordant between random and spouse pairs.

Assortative mating is likely to induce phenotypic and genetic structure between an individual and their spouse which complicates the interpretation of spousal comparisons in an aetiological context. A further consideration is that the joint participation of non-independent spouses in cohort studies could induce selection bias.

## Introduction

Human spouses are often highly similar for many traits ^1^. One mechanism underlying spousal similarities is that cohabitating spouses share a common environment during their relationship ^2^ which may act to increase phenotypic similarity, such as for behavioural (e.g., physical activity and alcohol use) or personality traits ^3; 4^. The shared adulthood environment between spouses has prompted the use of spouses in a variety of contexts in epidemiological and genetic research using a model that we refer to as the “within-spouse pair” (WSP) model. The WSP model involves modelling the similarities and differences of spouses, either by analysing the differences between each pair or by modelling spousal relationships as a covariate in a fixed-effect model. For example, previous studies have used the WSP model to estimate effects of the shared adulthood environment ^5–8^, while the WSP model has been proposed as an approach to reduce confounding bias in aetiological research with environmental confounders likely to be strongly correlated between spouses ^9^.

However, for many traits, there is evidence suggesting that phenotypic similarities do not substantially increase during a relationship ^10^. An alternative mechanism that induces spousal correlations is assortative mating – a phenomenon where humans are generally more likely to select a phenotypically similar ^3; 4; 11–15^ or, in some instances ^16^, dissimilar ^17; 18^ mate. For example, height and years in schooling are often fixed prior to partnership formation, suggesting that spousal similarities for these phenotypes reflect assortment rather than effects of the shared adulthood environment. Furthermore, geographical, ancestral and cultural factors often have strong influences on both phenotypic variation and partner selection patterns, illustrated by the ancestral similarities of spouses ^19^. Therefore, some degree of spousal phenotypic similarities is likely to be explained by spousal assortment on factors not typically defined as phenotypes, such as place of birth or religion.

Assortative mating complicates the interpretation of spousal analyses as it suggests that spousal similarities may not relate to the shared environment. Furthermore, the WSP model may be susceptible to collider bias. Collider bias occurs when conditioning on a variable which is influenced by two or more factors. This bias can induce spurious associations between these factors in the conditioned or selected sample. For example, associations between risk factors for a disease are likely to be distorted in samples of only diseased cases ^20; 21^. Similarly, spousal samples condition on spousal compatibility, a pairwise measure of how likely two individuals are to enter a relationship. If spousal compatibility is a function of phenotypic similarity across multiple phenotypes, then collider bias could potentially arise if we adjust for spousal pairing (**Figure 1**) ^20; 22^. Previous spousal studies have acknowledged assortative mating, but whether assortment could distort WSP comparisons has not been investigated extensively. For example, the possibility of collider bias has been little discussed. Therefore, we evaluated the use of spouses, (in e.g. the WSP model), for aetiological epidemiology.

**Figure 1:**
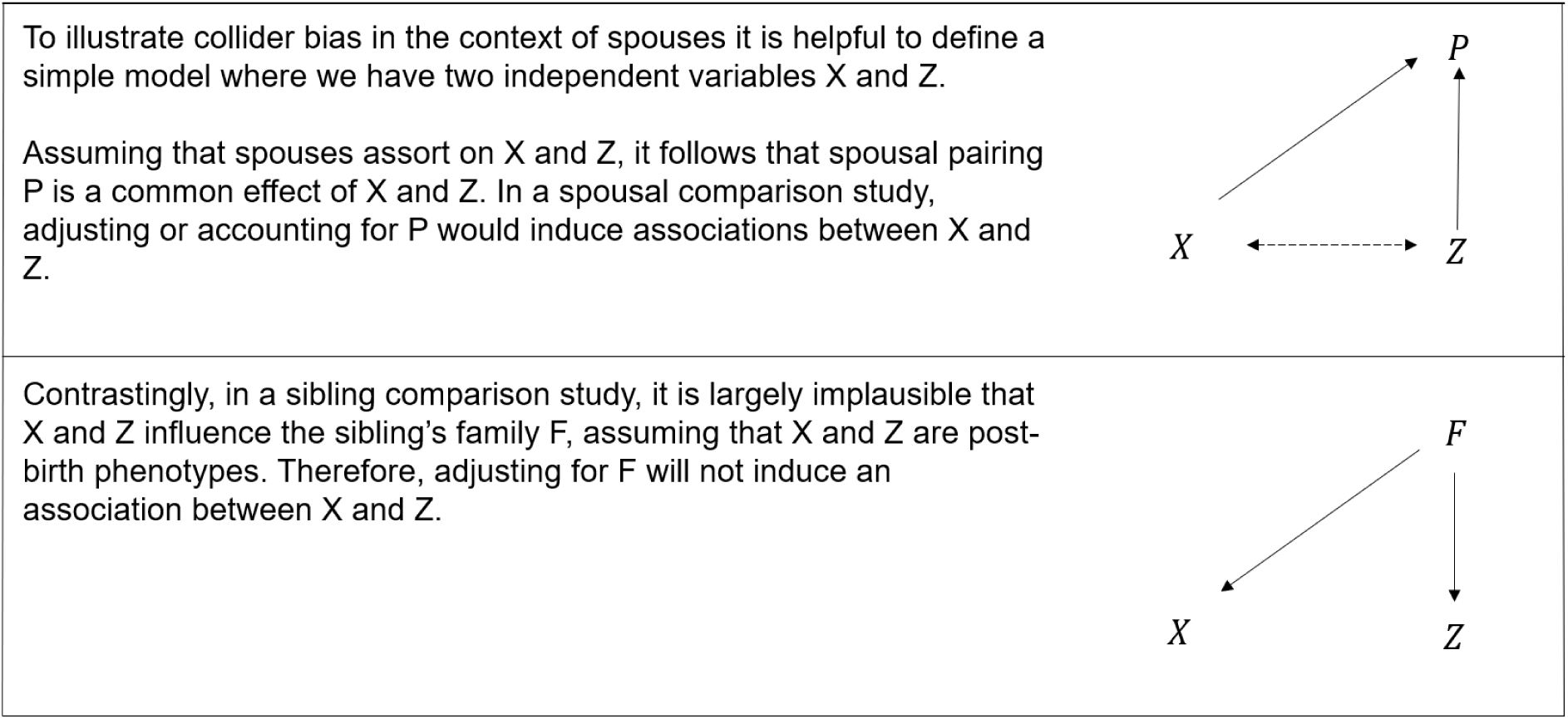
Potential for collider bias when comparing spouses and siblings.

First, we used directed acyclic graphs (DAGs) and simulated data to illustrate that the WSP model can be used to reduce confounding but is also susceptible to collider bias induced by assortative mating. Second, we used a WSP model applied to a sample of ~47,000 previously derived spouse-pairs from UK Biobank ^23^ to estimate the effects of increasing age on systolic blood pressure (SBP) and coronary artery disease (CAD). We generated and compared estimates from both spouse pairs and random non-assorted pairs, derived by reordering the spouse-pair sample. Third, we report estimates of spousal similarities, distinct from the WSP model, for phenotypic and genetic measures of height, educational attainment, body mass index, systolic blood pressure, coronary heart disease and alcohol consumption. Fourth, we used the WSP model to estimate effect sizes for genetic variants associated with these traits, compared with estimates from random pairs.

## Results

### Within-spouse pair model: assortative mating, spousal correlations and collider bias

Here, we present results from simulations evaluating the WSP model under assortative mating. In the first simulation model A, the relationship between an exposure and an outcome is confounded by an unmeasured factor. Spouses are positively correlated for the unmeasured confounder, either because of assortative mating or because of shared environmental factors during cohabitation. It follows that WSP estimates of the effect of the exposure on the outcome will be less biased (**Figure 2: panel A**).

In the second simulation model B, two independent exposures influence an outcome. Since assortment is influenced by the two exposures, it is a collider between them. It follows that the WSP estimates of the effect of either exposure on the outcome will be susceptible to collider bias dependent on the degree of assortment. However, if only one or neither exposure influences the outcome, the WSP estimates will not be affected (**Figure 2: panel B**).

**Figure 2:**
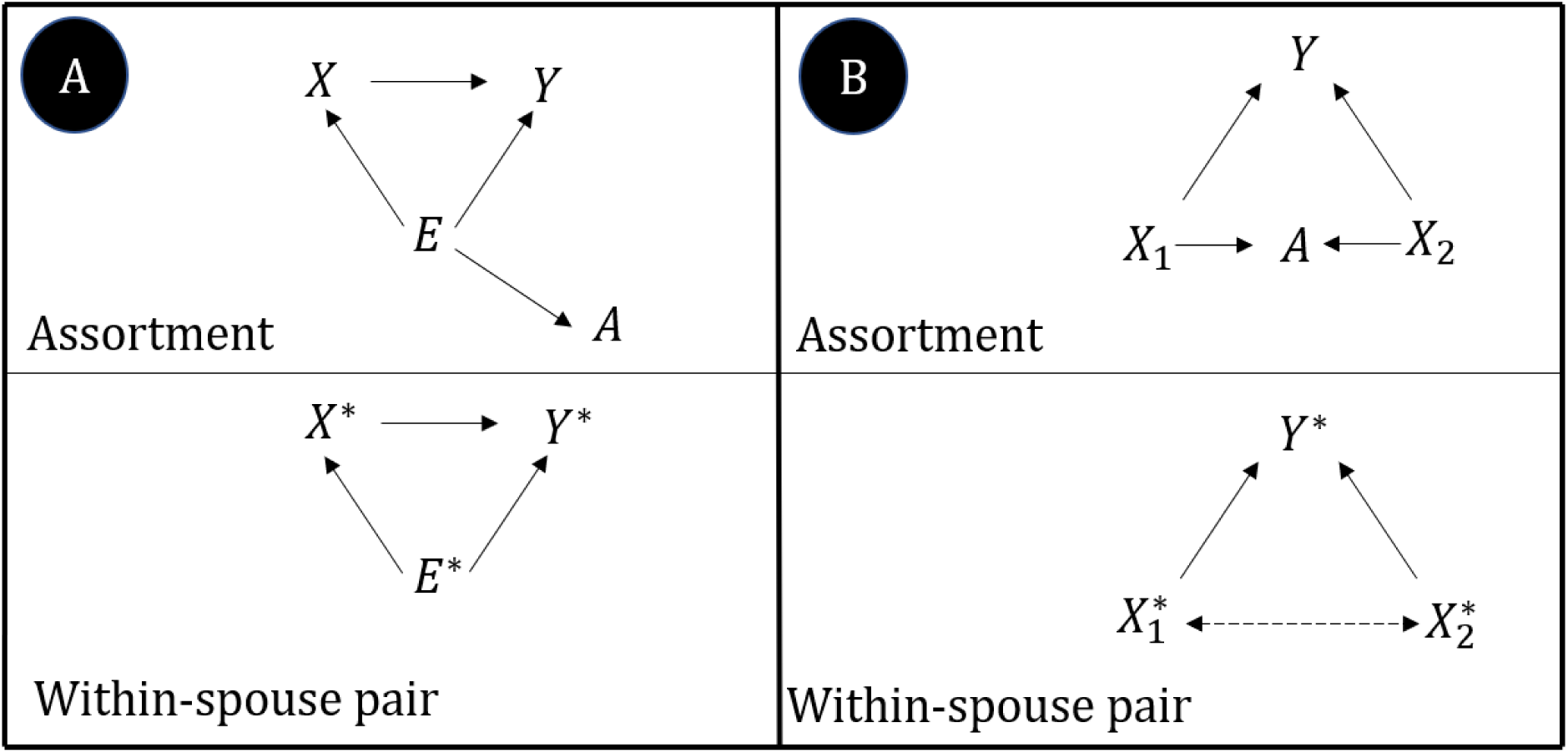
Simulated models of assortative mating, spousal correlations and collider bias.

The WSP design uses pairwise spousal differences (e.g. *X*_*M*1_ − *X*_*F*1_ & *Y*_*M*1_ − *Y*_*F*1_) in regression models, fitting each spouse pair as a single observation.

#### A) Within-spouse pair: spousal correlations for confounders

Exposure *X*; Outcome *Y*; Unmeasured confounder *E*; Spousal assortment *A*; WSP exposure *X*\* (*X*\* = *X*_*M*_ − *X*_*F*_); WSP outcome *Y*\* (*Y*\* = *Y*_*M*_ − *Y*_*F*_); WSP environmental confounder (the non-shared portion of the set of confounders) *E*\* (*E*\* = *E*_*M*_ − *E*_*F*_).

This figure illustrates the effect of an exposure on an outcome in the presence of an unmeasured confounder. Here, spousal pairing is determined by an assortment variable correlated with the confounder (indicated by A, a child of the confounder E). It follows that the value of spouses’ confounders will be correlated. In this example, a WSP model will reduce bias in the estimate of the effect of X on Y. In this figure we assume that spousal correlations for the confounder reflect assortment but in practice they could also relate to the shared spousal environment.

#### B) Within-spouse pair: assortative mating and collider bias

Exposures *X*_1_ *X*_2_; Outcome *Y*; Spousal assortment *A*; WSP exposures 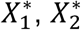; WSP outcome *Y*\* (*Y*\* = *Y*_*M*_ − *Y*_*F*_).

This figure illustrates the effect of an exposure on an outcome when two, otherwise independent exposures influence both the outcome and spousal assortment. It follows that associations will be present in the WSP model between the two exposures, which will distort the WSP estimated effect of the exposure on the outcome. We quantify the effect of potential collider bias in the WSP model at different levels of assortment on the two exposures.

Dashed lines indicate associations induced by spousal assortment.

### Simulations

We used simulated data to quantify the effect of the exposure on the outcome using the WSP model under two models. We generated 1,000 males (*M*1.. *M*1000) and 1,000 females (*F*1.. *F*1000). Male-female spouse pairs were determined using a spousal matching variable *A*, ordered such that *A*_*M*1_ ≥ *A*_*M*2_ ≥.. *A*_*M*1000_ and *A*_*F*1_ ≥ *A*_*F*2_ ≥.. *A*_*F*1000_, and pairs were then matched on the assortment variable, i.e. *A*_*M*1_ with *A*_*F*1_.

#### Model A: Within-spouse pair: spousal correlation for confounders

In this model we investigated the bias in WSP estimates of the effect of an exposure on an outcome if spouses assort on the measured confounder. The simulations demonstrate that the WSP estimate converges to the simulated unbiased estimate of 0.3 as the spousal correlation for the confounder tends to 1 (**Figure 3: panel A/Supplementary Table 1)**.

#### Model B: Within-spouse pair: assortative mating and collider bias

In this model, we evaluated the potential bias in WSP estimates when spousal assortment is a collider between two exposures. Simulations showed that the degree of bias in the effect estimate is a function of the degree of assortment on the two exposures with more bias when spouses strong assort on both traits. For example, under this model and using plausible assortment estimates for educational attainment (0.5) and height (0.2) ^24^, the expected bias would be around 13% when estimating the effect of education on a trait which is also influenced by height (**Figure 3: panel B/ Supplementary Table 2**).

**Figure 3:**
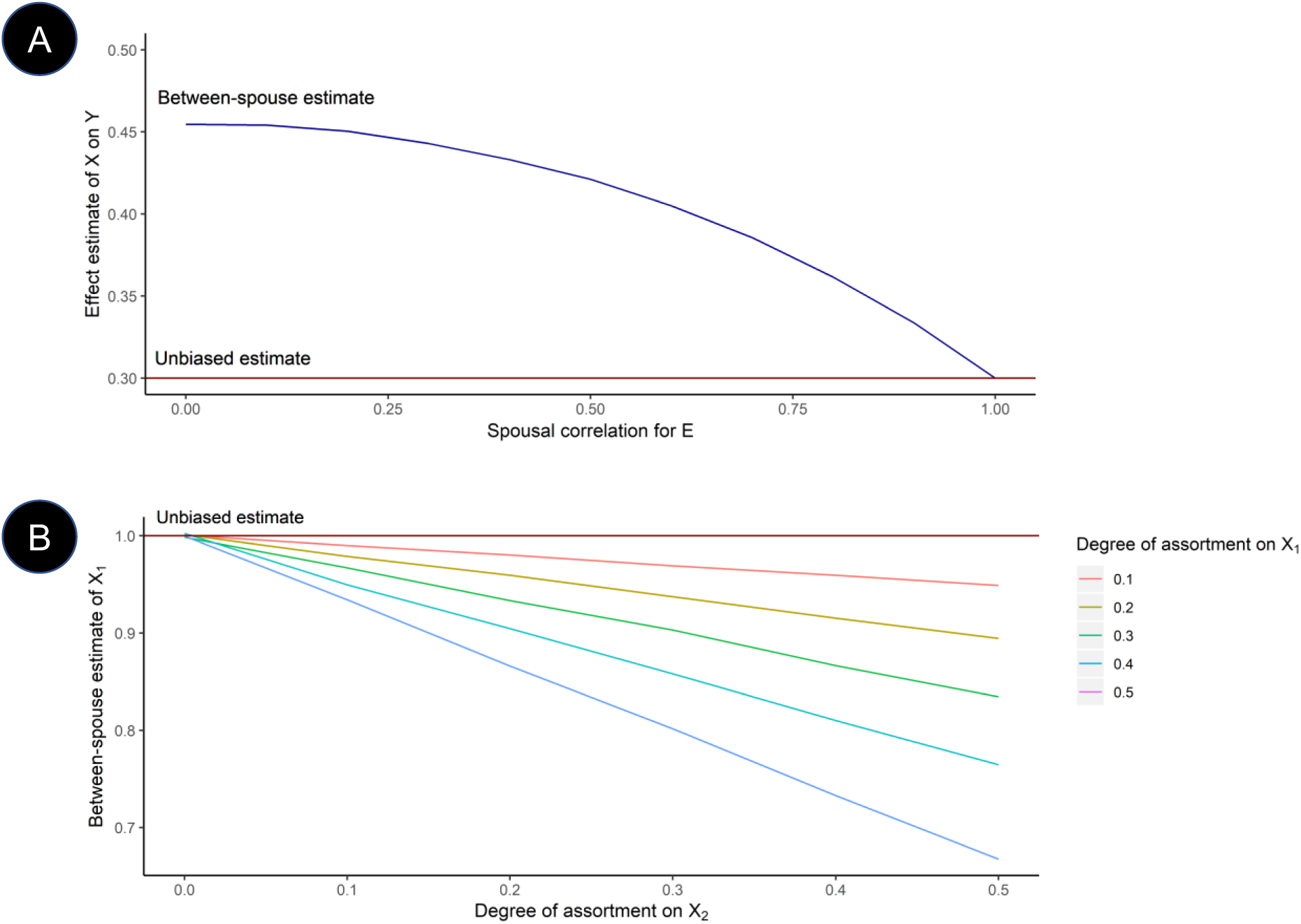
Simulation results for WSP models.

A – Simulations for Model A: Spousal correlations controlling for confounding.

As the strength of spousal assortment on the confounder (*E*) increases, the between-spouse association of *X*\* on *Y*\* unadjusted for *E* becomes less biased.

B – Simulations for Model B: Between-spouse: assortment and collider bias.

Spousal assortment can induce collider bias in WSP estimates. If spouses assort on two phenotypes *X*_1_ and *X*_2_ and affect outcome *Y*, then the association of *X*_1_ and *Y* is a biased estimate of the causal effect of *X*_1_ and *Y*. This bias monotonically increases in the degree of assortment on either *X*_1_ or *X*_2_.

### Empirical analyses using spouse pairs in UK Biobank

#### Within-spouse pair: age, SBP and CAD

We performed empirical analyses to evaluate the WSP design, using a sample of 47,435 spouse-pairs from UK Biobank, previously derived using household sharing information ^15^. The spouse-pair sample were broadly of European descent, as determined by principal component analysis, and had an average age of 59.5 years (on January 2010). Further characteristics of the sample for phenotypes of interest are contained in **Supplementary Table 3**.

As a positive control, we estimated the effects of increasing age on outcomes known to be related to age (CAD and SBP) in the spouse sample. For comparison, we calculated within-pair estimates using the same model in samples of non-assorted pairs. Pairwise age differences were found to be greater between random pairs which is evidence of assortative mating by age. We found consistent estimates of the effect of age on CAD and SBP between analyses conducted in the spouse and random pair samples (**Table 1**).

**Table 1:**
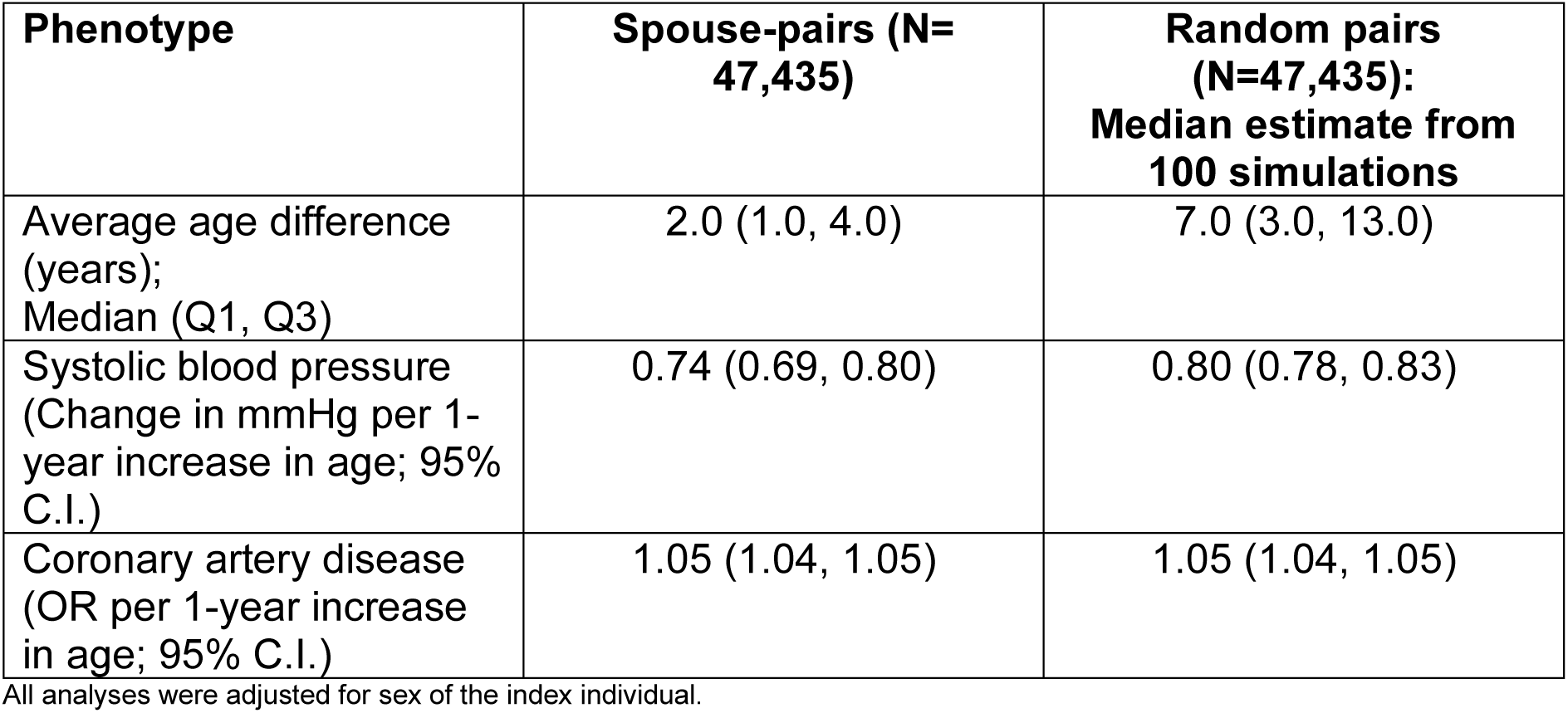
Increased age and age-related outcomes.

| Phenotype | Spouse-pairs (N=47,435) | Random pairs (N=47,435):<br>Median estimate from 100 simulations |
| --- | --- | --- |
| Average age difference (years);<br>Median (Q1, Q3) | 2.0 (1.0, 4.0) | 7.0 (3.0, 13.0) |
| Systolic blood pressure (Change in mmHg per 1-year increase in age; 95% C.I.) | 0.74 (0.69, 0.80) | 0.80 (0.78, 0.83) |
| Coronary artery disease (OR per 1-year increase in age; 95% C.I.) | 1.05 (1.04, 1.05) | 1.05 (1.04, 1.05) |

All analyses were adjusted for sex of the index individual.

#### Spousal phenotypic and genotypic correlations

We investigated the potential mechanisms influencing spousal phenotypic similarities using genetic data in UK Biobank ^13; 15; 24^. Spousal genetic associations (S_GG_) for genetic variants associated with a phenotype provide evidence for spousal assortment as genotype is fixed from birth and cannot be changed by the shared adulthood environment. Whereas, associations of an individual’s genotype and their partner’s phenotype (S_GP_) may relate to assortment or social genetic effects of an individual on their partner. Phenotypic correlations (S_PP_) between spouses may capture assortative mating, environmental effects or factors such as social homogamy, where individuals may assort on social factors which influence the exposure. Note that these analyses compare an individual to their spouse, distinct to the WSP model which compares spouse pairs to other spouse pairs.

We estimated spousal associations for height, educational attainment, body mass index, systolic blood pressure and coronary heart disease using both observed and genetic measures of the phenotypes. We scaled S_GP_ and S_GG_ associations to be on the same scale as the S_P×P_ estimates using an instrumental variable framework, Mendelian randomization for S_G×P_ and a similar approach for S_GG_. All phenotypes were concordant between spouses with some evidence of S_GP_ associations for height, education and body mass index. Notably, the S_PP_ associations were greater for all traits except for education where a stronger S_GP_ association was observed. Evidence of genotype-genotype correlations was evident only for height and educational attainment, with a substantial but imprecise scaled estimate for educational attainment (**Table 2**).

**Table 2:** Spousal phenotypic and genotypic associations.

| Phenotype | **$S_{PP}$ (95%CI):**<br>S.D. increase in individual phenotype per S.D. increase in spousal phenotype | **Scaled $S_{GP}$ (95%CI)**<br>S.D. increase in individual phenotype per S.D. increase in spousal GRS | **Scaled $S_{GG}$ (95%CI)**<br>S.D. increase in individual GRS per unit increase in spousal GRS |
| --- | --- | --- | --- |
| Height | 0.23 (0.22, 0.24) | 0.19 (0.17, 0.21) | 0.27 (0.12, 0.42) |
| Educational attainment | 0.46 (0.45, 0.46) | 0.71 (0.62, 0.80) | 2.00 (0.64, 3.36) |
| Body mass index | 0.24 (0.23, 0.25) | 0.15 (0.10, 0.21) | -0.27 (-0.95, 0.40) |
| Systolic blood pressure | 0.09 (0.08, 0.09) | 0.01 (-0.03, 0.04) | 0.06 (-0.27, 0.38) |
| Coronary heart disease | 0.02 (0.01, 0.03) | -0.04 (-0.13, 0.06) | -0.51 (-2.54, 1.53) |

All analyses were adjusted for sex and age

#### Within-spouse pair: genetic and phenotypic associations

We performed empirical analyses using genetic data to evaluate the WSP model in the context of evaluating genetic associations. Starting with Genome-wide Association Study (GWAS) data for height, SBP, body mass index, educational attainment and CAD, we extracted independent (r^2^ < 0.001) single nucleotide polymorphisms (SNPs) reaching genome-wide significance in previous GWAS (P<5×10^−8^) of traits of interest. We then estimated and compared SNP effect sizes when applying the WSP model to spouse and random pair samples. We found strong evidence of smaller effect sizes in the spouse sample for height (shrinkage: 23%; 95% CI 20%, 25%), educational attainment (shrinkage: 74%; 95% CI 66%, 81%) and body mass index (shrinkage: 23%; 95% CI 14%, 32%). There was some evidence of shrinkage for genetic variants associated with systolic blood pressure (shrinkage: 9%; 95% CI 1%, 17%), although this result would not pass adjustment for multiple testing, and limited evidence for coronary artery disease variants (**Table 3**).

**Table 3:** Genetic variant effect size comparison between spouse-pairs and random male-female pairs.

| Phenotype | Number of SNPs | Shrinkage in spouse-pair estimate compared to random pairs: % (95% C.I.) |
| --- | --- | --- |
| Height | 381 | 0.23 (0.20, 0.25) |
| Educational attainment | 69 | 0.74 (0.66, 0.81) |
| Body mass index | 68 | 0.23 (0.14, 0.32) |
| Coronary artery disease | 40 | 0.03 (-0.15, 0.22) |
| Systolic blood pressure | 190 | 0.09 (0.01, 0.17) |

For alcohol consumption, we considered only a single well-characterised variant in *ADH1B*. Each minor allele of the SNP was associated with an increased weekly alcohol consumption of 3.55 units (95% C.I. 3.03, 4.08) in the spouse sample compared to the median random-pair estimate of 4.98 units (95% C.I. 4.31, 5.64), an effect estimate shrinkage of 29% (95% C.I. 5%, 46%: heterogeneity P-value = 0.001).

## Discussion

In this study, we used directed acyclic graphs, simulations and empirical data to evaluate the use of the WSP model in aetiological epidemiology. We showed that the WSP model can account for unmeasured confounding if spouses are correlated for the confounder but that comparing assorted spouses can also induce collider bias. Empirically, we found that WSP effect estimates for age on coronary heart disease and systolic blood pressure were concordant between spouse and non-spouse pairs. Contrastingly, we found evidence that within-pair effect estimates for height, educational attainment, body mass index and alcohol consumption genetic loci are attenuated when comparing assorted spouses over non-spouse pairs.

There are several possible explanations for these attenuations. First, simulated data showed that if spouses assort on a confounder of the exposure and outcome, then the WSP association is a less biased estimate of the causal effect than a conventional model unadjusted for the confounder. An example of potential confounding is shared ancestry or place of birth, which is likely to be more correlated between spouses that for non-spouse pairs. Second, simulations also suggested that the WSP model applied to genetic associations may be susceptible to collider bias. Spousal assortment may induce biased associations between genetic and non-genetic influences on the trait of interest, with the degree of bias increasing with the strength of assortment, leading to the WSP model underestimating the true effect. The possibility that collider bias induced by assortment contributed to the shrinkage is supported by the greater effect size attenuations observed for traits with evidence of strong spousal assortment (height, educational attainment and alcohol consumption). Third, reduced phenotypic variation between spouses induced by assortment could lead to effect sizes attenuating towards the null because of the increased proportion of measurement error to variation. Fourth, the spousal environment could be influenced by individual’s genotypes leading to reduced spousal phenotypic differences. For example, if an individual has high genetic liability to increased alcohol consumption this could lead to their partner consuming similar amounts of alcohol independent of their genotype.

The WSP model is comparable to any covariate adjusted causal model where adjustment could both reduce confounding bias and induce collider bias ^25^. The merits of WSP estimates are dependent on their overall bias relative to the causal effect of interest. In empirical studies the size of this bias is likely to be difficult to quantify as the true causal effect will be unknown. In our positive control example of age on health outcomes, bias did not increase discernibly; we found little evidence that the within-pair effect estimates of age on CAD and SBP were distinct between spouse and non-spouse pair samples. These findings suggest that, in some instances, the WSP estimates are unlikely to be affected by collider bias, potentially when traits are not strongly assorted on as for CAD and SBP. Indeed, the phenotypic correlations for CAD and SBP (measured at study enrolment) are likely to reflect shared environmental factors rather than direct assortment.

A key implication is that spousal similarities and differences are not necessarily random or attributable solely to the shared adulthood environment. Amidst growing evidence that genetic epidemiological studies can be susceptible to bias from fine-scale population structure, dynastic effects and assortative mating ^26–35^, there is considerable interest in using genotype data from pedigrees to disentangle these effects and more accurately estimate trait heritability ^5–8; 26–28; 33; 34; 36–40^. Family designs such as the transmission disequilibrium test ^41^ and sibling comparisons are protected from many of these biases by random segregation at meiosis ^42; 43^. However, in contrast, inferences from spousal analyses are not as robust to bias, thus it is important to understand and model the assortment in spousal designs. A further implication is that assortative mating is likely to contribute to the phenotypic and genetic structure of epidemiological studies. Large studies such as the UK Biobank, frequently incidentally sample participants who are partnered with another study participant ^15^. These individuals will be non-independent. The consequences of jointly enrolled spouses are unclear but are one mechanism influencing study participation.

Previous studies have incorporated spousal relationships in variance component models as a measure of the shared environment during adulthood ^5–8^. Although researchers have acknowledged that assortative mating may induce bias in these models ^5^, the possibility of collider bias suggests that modelling spousal similarities without accounting for assortment could induce bias beyond the levels previously envisaged. It is difficult to discern whether WSP genetic associations are more accurate than conventional estimates; spousal assortment on ancestry may reduce bias, or collider bias induced by assortative mating may attenuate the associations. However, time spent in full-time education and height are unlikely to be affected by adult environment, providing evidence that effects are driven by spousal assortment. Our findings are consistent with recent work showing that assortative mating can induce bias in negative control studies ^44^, where paternal exposures are used as a control for effects of maternal exposures on offspring via the in-utero environment ^45^.

Our study has several important limitations. First, as described in our previous study ^15^, derived spouse-pairs were identified using household sharing information so may be susceptible to a degree of classification error with non-spouse pairs being incorrectly identified as spouses. Second, the mechanisms by which spouses jointly participate in UK Biobank may have induced selection bias into empirical analyses as these pairs could be more similar than pairs that did not jointly participate. Third, given that the exact mechanisms of assortment are not widely understood, our simulations and assumptions may not accurately capture the mechanisms underlying spousal assortment. In simulations we assumed that factors influencing assortment are independent across the population but in practice, factors influencing assortment are often correlated (e.g. height and education). Future research could use more complex simulations to evaluate models that can distinguish the effects of social homogamy, migration and measurement error. Fourth, it is important to note that educational attainment as defined by qualifications when study participants are aged over 40 will also capture individuals with degrees obtained during adulthood, suggesting that educational similarities could also plausibly relate to the shared adulthood environment.

To conclude, we provided evidence that WSP estimates for height, alcohol consumption, body mass index and educational attainment genetic variants are attenuated when compared to non-assorted pairs. Contrastingly, age-related associations were consistent between spouse and non-assorted pairs. Simulations suggested that a substantial proportion of these attenuations could be explained by assortative mating, emphasising the importance of considering assortative mating when using parental data in epidemiological and genetic studies.

## Methods

### Data sources

#### UK Biobank

##### Study description

UK Biobank is a large-scale prospective cohort study which sampled 503,325 individuals aged between 38-73 years at baseline, recruited between 2006 and 2010 from across the United Kingdom. The cohort has been described in detail previously ^23; 46^. For the purposes of this study, we used a subsample of the cohort of derived spouse-pairs, which has been described in detail in a previous study ^15^. In brief, 47,549 male-female pairs believed to be cohabitating spouses were identified from using household sharing information, including home coordinates (to the nearest km) and marital status, with closely related pairs identified and removed using a genetic relationship matrix.

##### Phenotype data

At baseline, the height of study participants was measured using a Seca 202 device at the assessment centre (field ID: 12144-0.0), body mass index was derived manually from measures of standing height and weight (field ID: 21001.0.0), systolic blood pressure was measured using an automated reading from an Omron Digital blood pressure monitor (field ID: 4080-0.0). Educational attainment was defined as in a previous study ^47^, using questionnaire data on qualifications to estimate the number of years spent in full-time education (field ID: 6138). Coronary artery disease cases were diagnosed using International Classification of Disease (10^th^ edition) (ICD10) and Operating Procedure System (OPS) codes from either hospital events (Hospital Episode Statistics) or underlying cause of death from the death register. The following ICD10 (I21, I22, I23, I24, I25, Z955) and OPS codes (K40-K46, K471, K49, K50, K75) ^48^ were used to classify diseased cases.

##### Genotyping

UK Biobank study participants (N= 488,377) were assayed using the UK BiLEVE Axiom™ Array by Affymetrix1 (N= 49,950) and the UK Biobank Axiom™ Array (N= 438,427). Directly genotyped variants were pre-phased using SHAPEIT3 ^49^ and then imputed using Impute4 using the UK10K ^50^, Haplotype Reference Consortium ^51^ and 1000 Genomes Phase 3 ^52^ reference panels. Post-imputation, data were available for approximately ~96 million genetic variants. More detail is contained in previous publications ^23; 53^.

#### Genome-wide Association Studies

Summary statistics from previous published GWAS, independent from UK Biobank, were used for information on SNPs associated with coronary artery disease ^54^, body mass index ^55^, educational attainment ^47^ and height ^56^.

Genome-wide summary data were not available for a recent systolic blood pressure GWAS ^57^, so we performed a GWAS of systolic blood pressure using UK Biobank. To remove sample overlap, we excluded the 47,539 spouse pairs from the analysis and used the remaining sample of 367,963 individuals of self-report European descent. A GWAS was conducted on this sample using a linear mixed model (LMM) association method as implemented in BOLT-LMM (v2.3)^58^. To model population structure in the sample we used 143,006 directly genotyped SNPs obtained after filtering on MAF > 0.01; genotyping rate > 0.015; Hardy-Weinberg equilibrium p-value < 0.0001 and LD pruning to an r2 threshold of 0.1 using PLINK v2.0 ^59^. We included the age and sex of participants as covariates in the model.

A set of Genome-wide significant SNPs were generated for each trait by LD clumping relevant summary statistics (P<5*×*10^−8^, r^2^<0.001, kb=10000) using the 1000 Genomes Phase 3 GBR samples ^52^ as the reference panel. For alcohol consumption, we used a missense variant (rs1229984) in *ADH1B* strongly associated with alcohol behaviour, as in a previous study ^15^.

### Theory of within-spouse pair comparisons

The phenotype *P* of individual *I* can be modelled as a function of independent factors; genetics *G*, the environment *E*, age, sex and a stochastic variance term ∈.

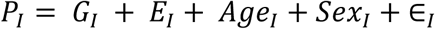

When considering male-female spouse pairs, we can decompose the influence of the environment *E* on *P* into effects of the shared environment between spouses *SE* (e.g. during cohabitation) and effects of the non-shared environment *NSE*. For example, for the male *M* and female *F* in pair *K*:

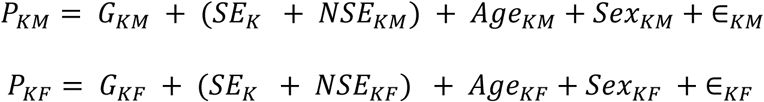

We then define the WSP model across spouse pairs as:

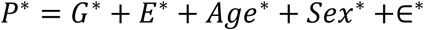

where the differences between the spouses for each factor are included in the model (e.g. for pair *K*,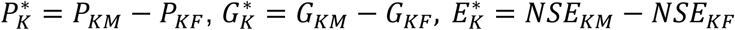). The shared environmental terms are by definition equal for men and women and drop out of the model.

For the WSP model to generate an unconfounded estimate of the causal effect of *G* on *P*, we require that the genetic and environmental difference terms in the between-spouse model are independent, i.e. *Corr*(*G*\*, *E*\*) = 0, an assumption that could be violated by several factors: e.g. population stratification bias or dynastic effects.

#### Random and non-random mating

Consider the between-spouse model applied to three distinct sets of pairs; a) a random set of males and females (non-spouses), b) spouse pairs under random mating (random spouses), and c) spouse-pairs under assortative mating (assorted spouses). In theory, the environmental differences between pairs would decrease with cohabitation and under assortment on environmental factors such as place of birth and socio-economic status:

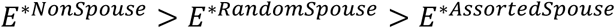

Note that as the environmental differences between pairs tends to zero (*E*\* → 0), the bias in the estimated association between *P* and *G* will also tend to zero (*bias*(*P*~*G*) → 0) even if *G*\* and *E*\* are correlated in the WSP model (*Corr*(*G*\*, *E*\*) ≠ 0) because the pair would be matched for the confounder, suggesting that comparing assorted pairs could reduce the effect of environmental biases.

We define the mechanism by which spouses assort as spousal compatibility *A*, a pairwise measure of how likely it is that two individuals enter a relationship. If several phenotypes influence assortment, then assortative mating can induce collider bias. For example, assortment on a phenotype influenced by genetic and environmental factors will induce spousal correlations in both genetic and environmental determinants of the phenotype, i.e. *Corr*(*G*_*KM*_, *G*_*KF*_) > 0 & *Corr*(*E*_*KM*_, *E*_*KF*_) > 0. It follows that in the WSP model, spousal genetic differences will be inversely associated with spousal environmental differences, i.e. *Corr*(*G*\*, *E*\*) < 0.

## Statistical methods

### Simulations

#### Model 1: Within-spouse pair: spousal correlation for confounders

In this model, in a sample of 1000 males and 1000 females, an exposure *X* influences an outcome *Y* but the relationship is confounded by life-course exposure to an environmental factor *E* which influences both *X* and *Y*. Spousal correlations for *E* are modelled by generating a spousal assortment measure *A*, correlated with *E* such that Corr(*E, A*)=*C*, that determines spousal pairing as follows. The spousal matching variable *A* is ordered such that *A*_*M*1_ ≥ *A*_*M*2_ ≥.. *A*_*M*1000_ and *A*_*F*1_ ≥ *A*_*F*2_ ≥.. *A*_*F*1000_, and male-female pairs are matched, i.e. *A*_*M*1_ with *A*_*F*1_. We evaluated whether spousal correlations for the confounder *E* influence the WSP estimates of *X* on *Y* by generating WSP estimates at a range of values of C (0.1, 0.2, 0.3, 0.4, 0.5, 0.6, 0.7, 0.8, 0.9).

#### Model 2: Within-spouse pair: assortative mating and collider bias

In this model, in a sample of 1000 males and 1000 females, individuals assort on two independent phenotypes *X*_1_ and *X*_2_, that also influence an outcome *0* such that *Y*~*X*_1_ + *X*_2_ +∈. Assortment is modelled by a spousal compatibility variable *A* which is correlated with *X*_1_ and *X*_2_ such that corr(*X*_1_, *A*)=*C*_1_ & corr(*X*_2_ *A*)=*C*_2_. The spousal compatibility variable is ordered for both males and females separately (*A*_*M*1_ ≥ *A*_*M*2_ ≥.. *A*_*M*1000_ and *A*_*F*1_ ≥ *A*_*F*2_ ≥.. *A*_*F*1000_), with pairs then matched on the compatibility variable, i.e. *A*_*M*1_ with *A*_*F*1_. In this context, we estimated the effect of *X*_1_ on *Y* using the WSP model.

We used simulations to evaluate the extent of collider bias with varying degrees of spousal assortment (*C*_1_/*C*_2_ = 0, 0.1, 0.2, 0.3, 0.4, 0.5). The WSP regression model is defined as *Y*\*~*X*_1_* where *Y*\* = *Y*_*KM*_ − *Y*_*KF*_ and 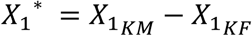 for each assorted pair.

### Empirical analysis in the UK Biobank

#### Within-spouse pair: age, SBP and CAD

In the sample of 47,435 spouse pairs, we defined the phenotypic differences between spouse pair *K* as follows:

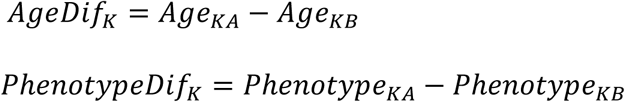

where *A* and *B* refers to the two individuals in the spouse pair. In the context of binary outcomes, the pair were rearranged so that the phenotypic difference could take the value of either 0 or 1 for the purposes of logistic regression, with other variables rearranged accordingly.

The WSP effect estimates of age on CAD and SBP were then estimated using the following regression model (linear or logistic dependent on the outcome of interest), including sex of the reference individual and the age difference between-spouses as covariates:

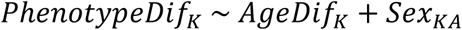

##### Random male-female pairs

As a point of comparison, we repeated the analysis using random male-female pairs to evaluate effects of spousal assortment. Using the same regression models described above, we repeated analyses using random male-female pairs. Starting with the 47,435 spouse-pairs, we generated 100 distinct datasets of random male-female pairs by rearranging couples at random. Across simulations, we reported the median effect size and standard error for each model.

#### Spousal phenotypic and genotypic similarities

We estimated associations between spouses for phenotypic and genotypic measures of height, educational attainment, BMI, SBP and CHD. Specifically, we estimated spousal phenotype-phenotype (S_PP_), genotype-phenotype (S_GP_) and genotype-genotype (S_GG_) associations for each trait.

We estimated S_PP_ associations using a regression of the observed trait in the index individual against the observed trait in their partner, including sex of the index individual and age of both spouses as covariates.

We estimated scaled S_GP_ associations using a Mendelian randomization framework ^43^, similar to a previous publication ^15^. To do this, we selected independent (r2<0.001 within 10,000kb) SNPs reaching genome-wide significance (p<5×10^−08^), which we used to construct a genetic risk score (GRS) using allele counts and effect sizes from the GWAS summary data. We then tested whether GRS for each trait were associated with a) phenotypes in the index individual (adjusting for sex, age and the first 10 PCs of index individual) and b) with the trait in the partner (adjusting for sex of index individual and age/ first 10 PCs of both spouses) using linear models in R. As in Mendelian randomization, the scaled S_GP_ estimate is defined as the ratio of these estimates (estimate b divided by estimate a) and was calculated using the seemingly unrelated regression (SUR) package in R which accounts for correlation between standard errors.

We used a similar framework to above to estimate the scaled S_GG_ associations. Using the GRS described above, we tested whether GRS for each trait were c) associated with the same GRS in their partner, including sex of the index individual and age of both spouses as covariates. By analogy with Mendelian randomization, the S_GG_ estimate can be scaled by dividing the genetic correlation estimate (estimate c) by the association of the GRS with phenotype in the same individual (estimate a) squared. Again, we calculated the scaled S_GG_ associations using SUR in R. Relevant mathematical equations are contained in the supplementary material of a previous manuscript ^15^.

#### Within-spouse pair: genetic and phenotypic differences

We defined the genotype difference for spouse pair *K* for each variant of interest as:

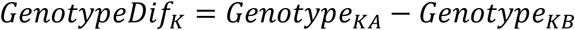

The WSP effect estimates of each genetic variant on the relevant phenotype of interest (height, body mass index, systolic blood pressure, educational attainment, coronary artery disease or alcohol consumption) were estimated using the following regression model (again linear or logistic), including sex of the reference individual and age difference between the spouses as covariates:

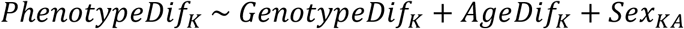

##### Comparing between-spouse and between-random pair estimates across genetic variants

As above, we calculated the associations using the WSP model using the actual spouses and then repeated analyses using 100 distinct datasets of random male-female pairs. We then took the ratio of effect estimates between the spouse sample estimate and the median effect estimate from the 100 random-pair estimates, with standard errors of ratios calculated using the delta method. A fixed effects model using the metafor package ^60^ in R was then used to meta-analyse the ratios across all SNPs for each trait of interest assuming independence of SNPs. A shrinkage estimate was then generated by subtracting the meta-analysis result and confidence interval from 1.

We removed outlying variants for each trait to prevent convergence problems in the meta-analysis. Outliers were defined as variants where the ratio of effect sizes between the spouse and random-pair models were more than five times the interquartile range away from the mean ratio. In practice we removed two outlying variants for educational attainment.

As we investigated only a single genetic variant for alcohol consumption, we were unable to investigate a trend across genetic variants. Instead we tested for a difference between two means for the WSP and median random-pair estimate ^61^.

## Data and code availability

Relevant code for simulations is available at the following repository https://github.com/LaurenceHowe/Between-spouse. A list of derived spouse-pairs is available on request from UK Biobank.

## Acknowledgements

The Medical Research Council (MRC) and the University of Bristol support the MRC Integrative Epidemiology Unit [MC_UU_12013/1, MC_UU_12013/9, MC_UU_00011/1]. The Economics and Social Research Council (ESRC) support NMD via a Future Research Leaders grant [ES/N000757/1]. No funding body has influenced data collection, analysis or its interpretation. This work is part of a project entitled ‘social and economic consequences of health: causal inference methods and longitudinal, intergenerational data’, which is part of the Health Foundation’s Efficiency Research Programme (Award 807293). The Health Foundation is an independent charity committed to bringing about better health and health care for people in the UK. This publication is the work of the authors, who serve as the guarantors for the contents of this paper.

