## Supplementary Material for "Assortative mating and within-spouse pair comparisons"

**Supplementary Table 1** Model 1: Spousal correlations controlling for confounding

| **Degree of spousal correlation for the confounder** $\boldsymbol{E}$ | **WSP effect estimate of exposure** $\boldsymbol{X}$ **on outcome** $\boldsymbol{Y}$ **(unadjusted for** $\boldsymbol{E}$**):**  Simulation mean |
| --- | --- |
| True unconfounded estimate | 0.30 |
| 0 | 0.46 |
| 0.1 | 0.45 |
| 0.2 | 0.45 |
| 0.3 | 0.44 |
| 0.4 | 0.43 |
| 0.5 | 0.42 |
| 0.6 | 0.41 |
| 0.7 | 0.39 |
| 0.8 | 0.36 |
| 0.9 | 0.33 |

**Supplementary Table 2** Model 2: Assortment and collider bias

| **Degree of assortative mating on** $\boldsymbol{X}_{\boldsymbol{1}}$ | **Degree of assortative mating on** $\boldsymbol{X}_{\boldsymbol{2}}$ | **WSP effect estimate of exposure** $\boldsymbol{X}_{\boldsymbol{1}}$ **on outcome** $\boldsymbol{Y}$**:**  Simulation mean as proportion of true effect size |
| --- | --- | --- |
| 0 | (0, 0.1, 0.2, 0.3, 0.4, 0.5) | 1.00 |
| 0.1 | 0.1  0.2  0.3  0.4  0.5 | 0.99  0.98  0.97  0.96  0.95 |
| 0.2 | 0.1  0.2  0.3  0.4  0.5 | 0.98  0.96  0.94  0.92  0.90 |
| 0.3 | 0.1  0.2  0.3  0.4  0.5 | 0.97  0.93  0.90  0.87  0.83 |
| 0.4 | 0.1  0.2  0.3  0.4  0.5 | 0.95  0.91  0.86  0.81  0.76 |
| 0.5 | 0.1  0.2  0.3  0.4  0.5 | 0.93  0.87  0.80  0.73  0.67 |

**Supplementary Table 3** Characteristics of the spouse sample (N$\leq$94,870)

| **Phenotype** | **Mean (SD)** |
| --- | --- |
| Age on 1^st^ January 2010 (years) | 59.5 (7.4) |
| Height (cm) | 169.2 (9.3) |
| Educational attainment (years in full-time schooling) | 14.1 (2.3) |
| Systolic blood pressure (mmHg) | 141.3 (19.5) |
| Body mass index (kg/m^2^) | 27.4 (4.5) |
| Coronary artery disease: cases (% of sample)^1^ | 4664 (4.9%) |

^1 Dichotomous trait so number of cases and % presented^
